# Epiphytic Microbiome of Grapes Berries Varies Between Phenological Timepoints, Growing Seasons and Regions

**DOI:** 10.1101/2019.12.20.884502

**Authors:** Megan E. Hall, Isabelle O’Bryon, Wayne F. Wilcox, Michael V. Osier, Lance Cadle-Davidson

## Abstract

Extensive research into the microbial ecology of grapes near harvest, with a primary focus on yeasts, has improved our understanding of some components of variation that influence grapevine terroir. Metagenomic tools enable a broader exploration of the plant microbiome and components of variability due to such factors as year, location, management, and phenological stage. In 2014, to characterize the microbial changes of the grape surface over the course of the growing season in the Finger Lakes region of New York, we examined the epiphytic microbiome of grapes at five key phenological stages: pea-sized, bunch closure, Veraison, 15 Brix and harvest. This experiment was repeated in two subsequent years in the Finger Lakes, New York in 2015, and in Tasmania, Australia in 2016, to examine variability of regional terroir. We found significant shifts in taxa presence and relative taxa abundance between phenological timepoints, and determined that the epiphytic microbiome differed significantly not just between regions but also within a single region from one year to the next. These findings call into question the role of the phytobiome in the expression of terroir, as the phytobiome is dynamically responding to its environment, within and between years and locations. On the berry surface in particular, these dynamics are complicated by weather and management. Understanding that the grape surface microbiome is consistently changing may influence how we manage the berry epiphytic microbiome, potentially affecting disease management and vinification decisions.

## Introduction

Recent research into the microbiota of grapes examined the microbial communities constituting a particular microbial terroir through sampling of grape at harvest or in the grape must after harvest. Microbial sampling has been examined in vineyards determined to have the same terroir (Setati et al., 2012), and native microbial populations examined across regions (Martiny et al., 2006), but the changes in one region across an entire season and between two regions in multiple years has not been explored. While the microbial populations on grapes immediately before harvest has been extensively investigated (Brysch-Herzberg and Seidel, 2015; Combina et al., 2005; Drożdż et al., 2015; Garijo et al., 2011; Garofalo et al., 2016; Hall et al., 2019a; Jara et al., 2016; Martini et al., 1996; Parish and Carroll, 1985; Raspor et al., 2006; Rosini et al., 1982; Sabate et al., 2002; Setati et al., 2015; Yanagida et al., 1992) and while some researchers have investigated changes in microbial populations for the last few weeks before harvest (Garijo et al., 2011; Renouf and Lonvaud-Funel, 2007), fluctuations of microbial populations from the very beginning of berry development until harvest have not been investigated. Recently, a distinction has been made between the epiphytic and endophytic microbiota of grapes (Hall and Wilcox, 2018), which brings up new questions about the fluctuations of these separate microbiotas over the course of the growing season. Little is known about how the epiphytic microbiome of grapes changes throughout the growing season, partially because of the challenge of extracting sufficient DNA from young grape, and solely from the surface, yet understanding these fluctuations could influence management decisions. Frequent spray applications may be influencing the microbiome of the grape surface, which in turn may affect how the grape responds to both beneficial and pathogenic microbes. The microbial communities that are brought into the winery after harvest are not static, and the dynamics of the system could also affect fermentation in the winery.

## Materials and Methods

### Sample Collection

In 2014 and 2015, grapes were collected from two commercial vineyards, one of *Vitis vinifera* cv. Riesling and one of cv. Pinot Gris and one research vineyard of *Vitis* interspecific hybrid cv. Vignoles, all in the Finger Lakes region of New York. One additional commercial vineyard was added in 2015 with a planting of cv. Vignoles, also in the Finger Lakes region. In 2016, grapes were collected from five commercial vineyard blocks, one of *V. vinifera* cv. Sauvignon Blanc, and four *V. vinifera* cv. Riesling in Tasmania, Australia. To address fluctuations in microbial populations both within a vineyard and on an individual cluster, as articulated by Barata et al. (2012), we sampled individual berries, as opposed to whole clusters, and at varying locations in the vineyard. In every vineyard block, 12 panels were randomly selected and one cluster was randomly selected at the following phenological time points: pea-sized berries, bunch closure, Veraison, 15 Brix and harvest, for a total of 12 samples per time point per vineyard. In 2014, 180 samples were collected, in 2015, 240 samples and in 2016, 300 samples. The first three sampling points were determined by visually assessing the clusters in the 12 panels, and harvesting samples when 50% of the berries on a randomly selected cluster were determined to be at that particular phenological stage. For the sampling point of 15 Brix, 20 berries were selected randomly from each of three individual rows, and samples were collected when the juice averaged 15 Brix by refractometer. The harvest date for all years was determined when the fruit reached an average of 23 to 24 Brix. 20 berries were selected randomly from each of three individual rows, and samples were collected when the juice averaged 23-24 Brix by refractometer. Each randomly selected cluster was marked with flagging tape so as not to be sampled again at a future sampling point, which ensured that any changes to the cluster architecture or surface microbiota caused by sampling would not influence other samples. Three randomly selected berries, located at the tip of the cluster, the anterior side and posterior side, were cut from each cluster above the pedicel using scissors that were immersed in 95% ethanol between samples, and dropped directly into 50 mL Falcon tubes containing 5 mL of 10% w/v NaCl in TE buffer (10mM Tris-HCl+1mM EDTA, ph 8.0). The caps were screwed back on each tube immediately, and were placed in a Styrofoam cooler containing an ice pack until they were transported to the laboratory.

### DNA extraction

In the laboratory, 500 μl of 10% SDS was added to the Falcon tube containing the grape berry and TE-NaCl solution, vortexed for 5 seconds and left at room temperature for 15 minutes. A freeze-thaw sequence consisting of 30 minutes in a −80 C freezer and 5 minutes in 60 C water bath was repeated three times to lyse the fungal and bacterial cells. 750 μl of the solution was transferred to a centrifuge tube, along with 750 μl ice-cold isopropanol. The solution was centrifuged for 10 minutes at 9600 *g*. The supernatant was carefully removed from the tube, 500 μl ice-cold 95% ethanol was added, and the tube was again centrifuged at 9600 *g* for 1 minute (Hall et. al 2019b). The pellet was re-suspended in 100 μl TE buffer. The DNA was then stored at 4 C until further use.

### Amplification and Sequencing

Genomic DNA was sent to the Cornell University DNA Sequencing facility in Ithaca, NY for Illumina 250-bp-paired-end sequencing on the Illumina MiSeq machine. For each sample, two separate runs were performed. To amplify the V4 domain of bacterial 16S rRNA genes, primers F515 (5′*NNNNNNNN*GTGTGCCAGCMGCCGCGGTAA–3′) and R806 (5′–GGACTACHVGGGTWTCTAAT–3′) and for fungal internal transcribed spacer (ITS) 1 loci were amplified using primers BITS (5′-*NNNNNNNN*CTACCTGCGGARGGATCA–3′) and B58S3 (5′–GAGATCCRTTGYTRAAAGTT–3′) (Bokulich et al., 2014). Both forward primers were modified to contain a unique 8-bp barcode, highlighted in the italicized N-sections above.

### Data Analysis

Quality filtering, read processing, and OTU assignment was conducted in Qiime 1.9.1 (Caporaso et al., 2010b). Sequences were trimmed once there were three consecutive bases with PHRED scores less than 20. Sequences less than 100nt were discarded. Open and closed reference OTU-picking methods used uclust and a pairwise identity of 97% (Edgar, 2010). Alignment to greengenes 13_5 was done using PyNAST and alignment to UNITE 7_97 was conducted using the BLAST alignment method (Altschul et al., 1990; Caporaso et al., 2010a; DeSantis et al., 2006; Kõljalg et al., 2013). Rare OTUs were filtered if they had less than 0.0001% of the total abundance from within the biom file. Biom files were converted into spf files using the biom_to_stamp.py script provided by STAMP. The original mapping file and the spf file were read into STAMP, and an ANOVA test was done using the Tukey-Kramer method set to 0.95 and a p value filter of 0.05. The percentage of each taxon in each sample was calculated. The mean of the percentages for each taxon within each treatment was calculated and plotted in R. Organisms that could not be identified to the family level were excluded from the analysis. Heatmaps were made in R v.3.3.2 using the pheatmap package (R Core Team 2013, Kolde 2012). The colors represent the log of the relative mean frequency for each taxon. If a taxon was not seen in a given group the value was assigned to the lowest value in the matrix. Hierarchical clustering was done using the complete method, the rows were clustered using the Euclidean method, and the columns were clustered using the Manhattan method.

## Results

The sampling strategy focused on isolation of DNA from the epiphytic microbiome of three grape berries per sample. Examining all years at once, the ability to detect taxa from this small biomass generally increased over the course of the growing season (fungal samples: P=0.76; bacterial samples: P=0.75), with the highest percentage reads coming from Veraison and later (Table 1). The number of OTUs detected in both fungal and bacterial communities varied significantly among developmental stages, location, and year. In 2014, the least number of fungal and bacterial OTUs were detected than in any other year of the study (20 and 15, respectively). *Mucor* spp. represented 33% of the OTUs found at the bunch closure stage, 45% of those found at Veraison, but none at 15 Brix and then again 19% at harvest. One species of *Aspergillus (A. piperis)* was detected only at pea-sized berries and bunch closure, a different species (*A. flavus)* was detected only at Veraison through harvest. *Botrytis caroliniana* was detected for the first time at Veraison and at an increasing rate until harvest, where it occupied nearly half of the read. Two yeast genera (*Candida* and *Talaromyces*) were detected at only a single timepoint. Four species of *Penicillium* appeared at three of the five time points sampled, but not in consecutive order and not in any increasing or decreasing numbers throughout the growing season (Table 2). Within the bacterial reads, several species were found in every sampled time point. For instance, *Acinetobacter rhizosphaerae* represented nearly 85% of the reads at pea-sized berries, and remained at approximately 30% of the total OTUs for the rest of the growing season, whereas *Anoxybacillus kestanbolensis* was found in 4 – 11% of every sampled timepoint. *Pseudomonas balearica* was consistently found in each timepoint yet varied significantly, reaching its peak at 61% of the OTUs at Veraison (Table 3).

**Table 1.**
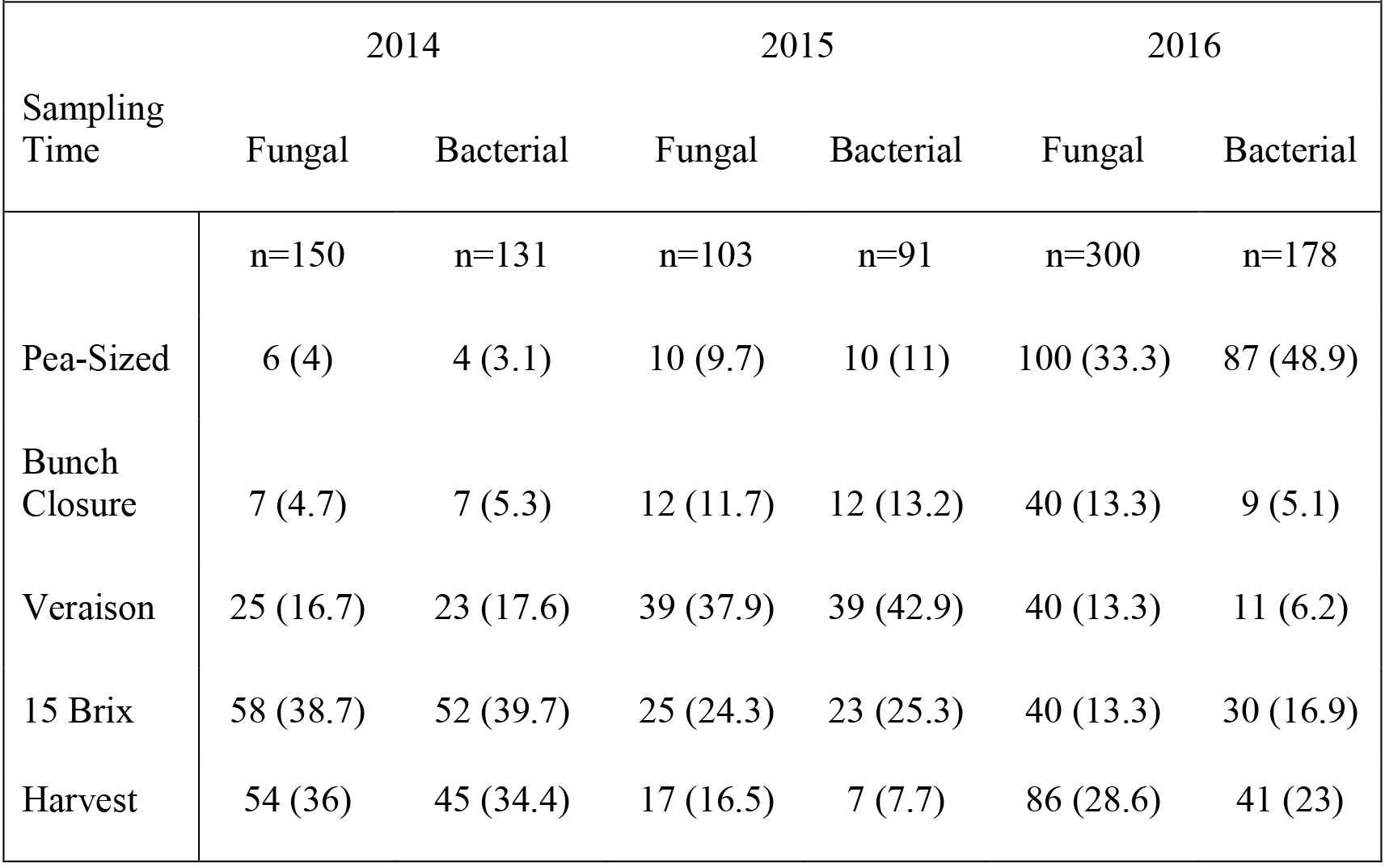
Number of samples (percent) passing quality filtering and OTU assignment by phenology, year, and Kingdom.

| Sampling Time | 2014 |  | 2015 |  | 2016 |  |
| --- | --- | --- | --- | --- | --- | --- |
|  | Fungal | Bacterial | Fungal | Bacterial | Fungal | Bacterial |
|  | n=150 | n=131 | n=103 | n=91 | n=300 | n=178 |
| Pea-Sized | 6 (4) | 4 (3.1) | 10 (9.7) | 10 (11) | 100 (33.3) | 87 (48.9) |
| Bunch Closure | 7 (4.7) | 7 (5.3) | 12 (11.7) | 12 (13.2) | 40 (13.3) | 9 (5.1) |
| Veraison | 25 (16.7) | 23 (17.6) | 39 (37.9) | 39 (42.9) | 40 (13.3) | 11 (6.2) |
| 15 Brix | 58 (38.7) | 52 (39.7) | 25 (24.3) | 23 (25.3) | 40 (13.3) | 30 (16.9) |
| Harvest | 54 (36) | 45 (34.4) | 17 (16.5) | 7 (7.7) | 86 (28.6) | 41 (23) |

**Table 2.**
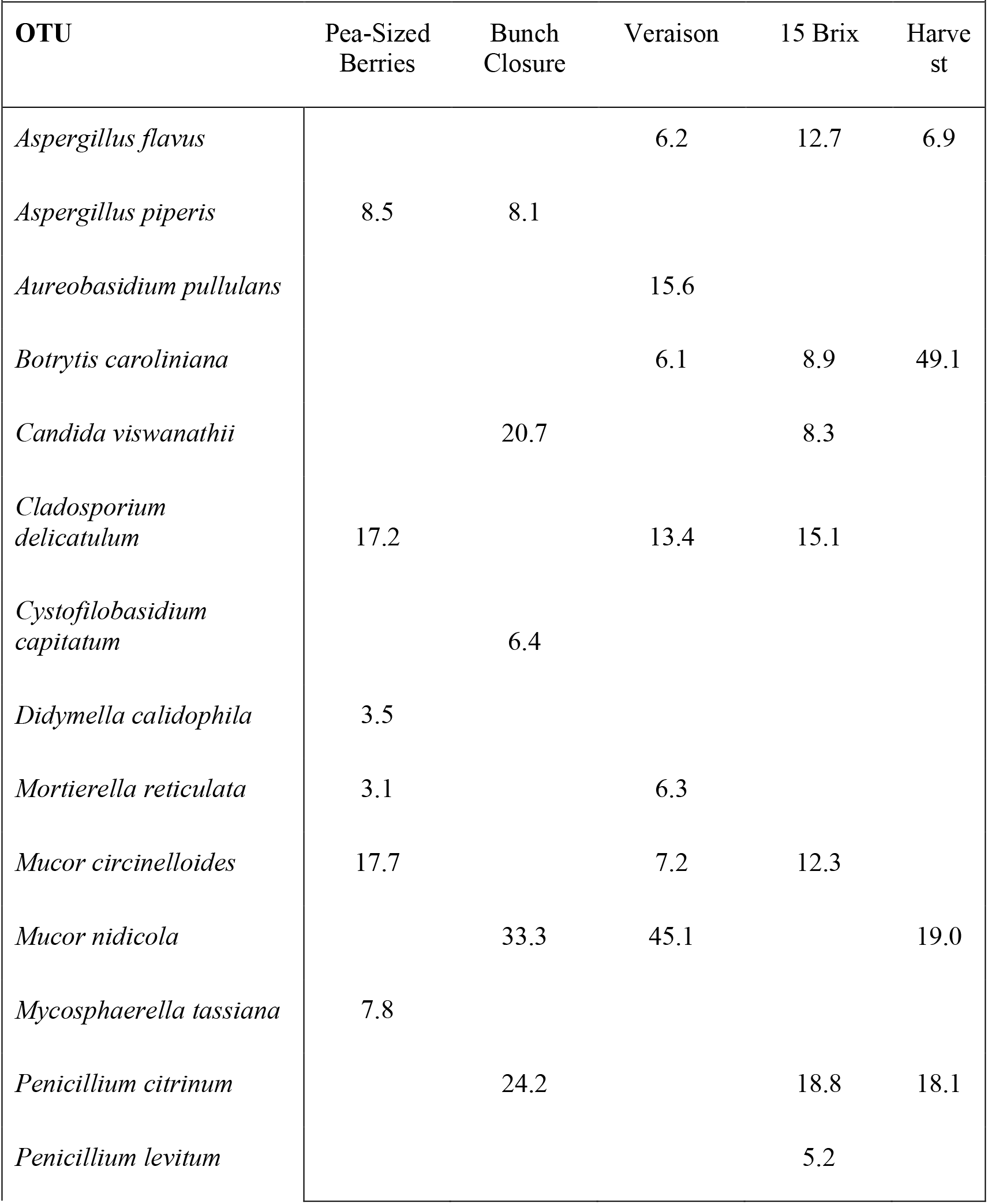
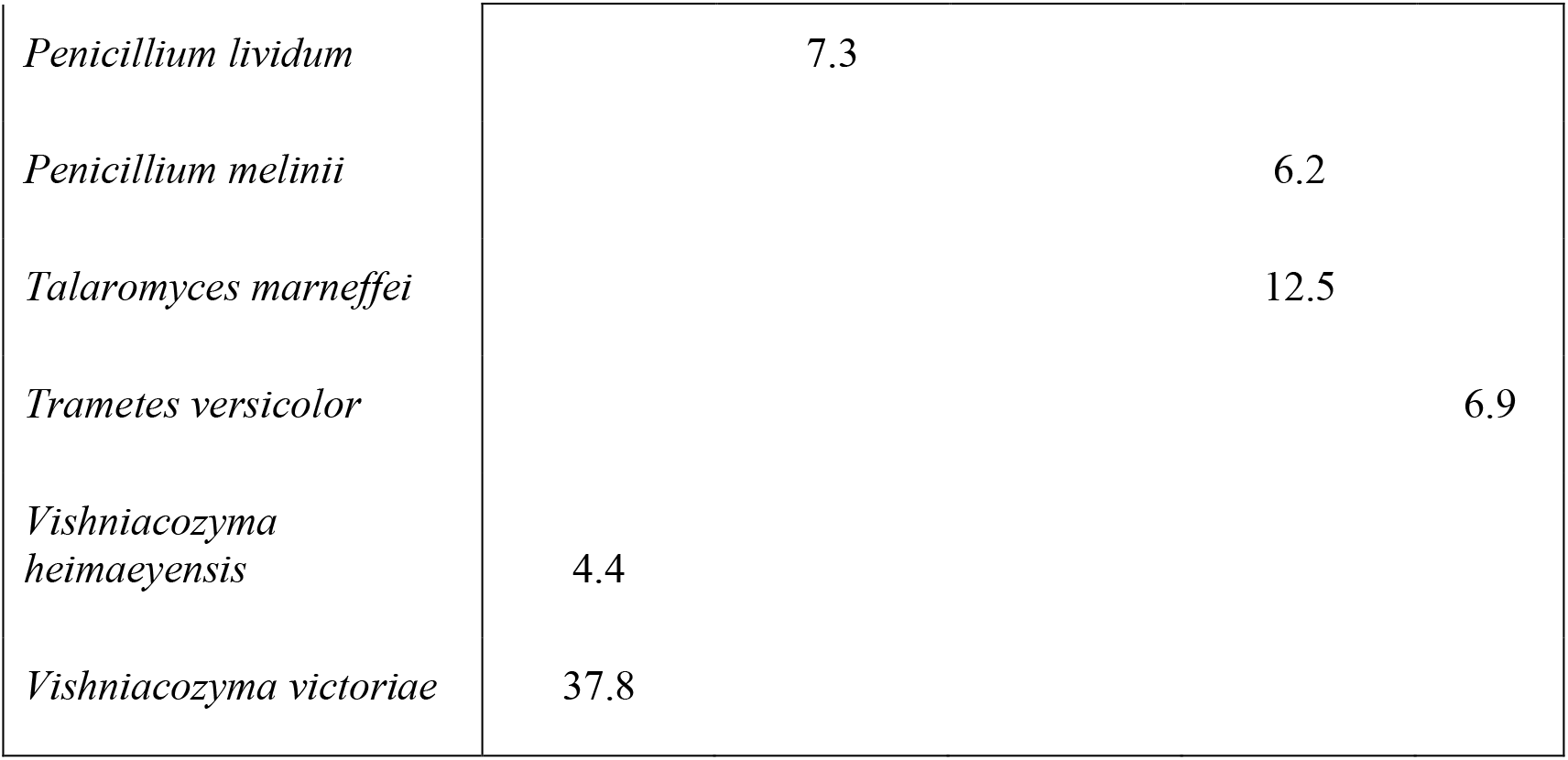
In 2014 Finger Lakes, New York, the relative mean frequency (%) of reads for each fungal OTU across three vineyards at five phenological stages. Sample numbers per stage are presented in Table 1.

**Table 3.**
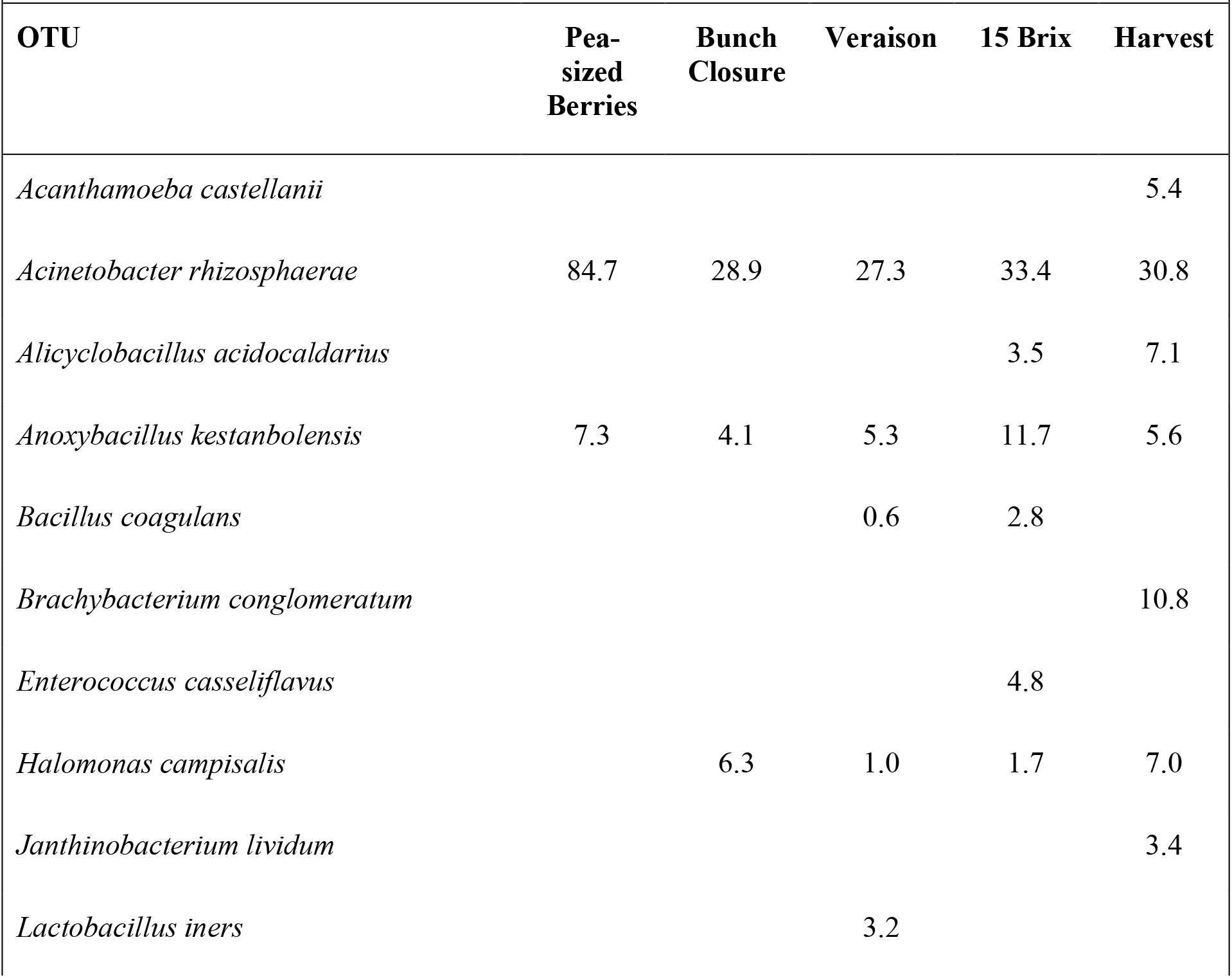
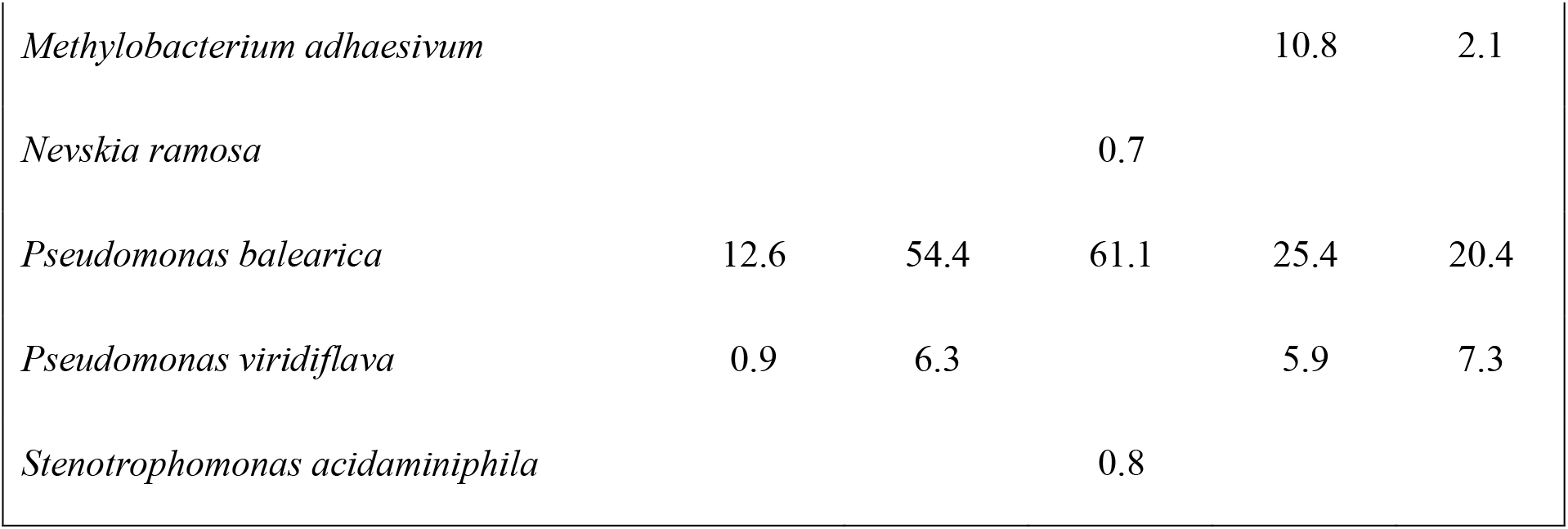
In 2014 Finger Lakes, New York, the relative mean frequency (%) of reads for each bacterial OTU across three vineyards at five phenological stages. Sample numbers per stage are presented in Table 1.

|  |  |  |  |  |  |
| --- | --- | --- | --- | --- | --- |
| **OTU** | **Pea-sized Berries** | **Bunch Closure** | **Veraison** | **15 Brix** | **Harvest** |
| *Acanthamoeba castellanii* |  |  |  |  | 5.4 |
| *Acinetobacter rhizosphaerae* | 84.7 | 28.9 | 27.3 | 33.4 | 30.8 |
| *Alicyclobacillus acidocaldarius* |  |  |  | 3.5 | 7.1 |
| *Anoxybacillus kestanbolensis* | 7.3 | 4.1 | 5.3 | 11.7 | 5.6 |
| *Bacillus coagulans* |  |  | 0.6 | 2.8 |  |
| *Brachybacterium conglomeratum* |  |  |  |  | 10.8 |
| *Enterococcus casseliflavus* |  |  |  | 4.8 |  |
| *Halomonas campisalis* |  | 6.3 | 1.0 | 1.7 | 7.0 |
| *Janthinobacterium lividum* |  |  |  |  | 3.4 |
| *Lactobacillus iners* |  |  | 3.2 |  |  |
| *Methylobacterium adhaesivum* |  |  |  | 10.8 | 2.1 |
| *Nevskia ramosa* |  |  | 0.7 |  |  |
| *Pseudomonas balearica* | 12.6 | 54.4 | 61.1 | 25.4 | 20.4 |
| *Pseudomonas viridiflava* | 0.9 | 6.3 |  | 5.9 | 7.3 |
| *Stenotrophomonas acidaminiphila* |  |  | 0.8 |  |  |

The 2015 season was significantly different from the 2014 season, in that more than double the number of fungal OTUs were detected (44) and almost four times the number of bacterial OTUs (56). At pea-sized berries no single OTU represented more than 18% of the reads. Many more yeast were present in 2015 than in 2014, such as *Metchnikowia* spp., *Pichia* spp., and *Sporobolomyces* spp. *Pichia kluyveri* oscillated at each developmental stage, representing 5.7% of the total reads at pea-sized berries, just 0.5% of reads at bunch closure, 47% at Veraison, only 5% at 15 Brix, and no reads at harvest. A similar pattern occurred with *Sporobolomyces ruberrimus*, with lows and highs ranging from 0.73% of OTUs at Veraison to over 87% at the very next phenological stage, 15 Brix. At harvest, three species dominated as *Botrytis caroliniana* represented 26.5% of the reads, *Coriolopsis gallica* represented 46% and *Metchnikowia pulcherrima* represented 26.5% (Table 4). Like the fungal reads, many of the bacterial OTUs throughout the 2015 growing season represented less than 1% of the total reads. Members of the family Burkholderiaceae dominated the reads, representing 39% of reads at pea-sized berries, 3% at bunch closure, 73% at Veraison, 86% at 15 Brix, and 65% at harvest. At harvest, Acetobacteraceae represented 5% of reads, *Gluconobacter* 4%, and *Gluconacetobacter* 4% (Table 5).

**Table 4.**
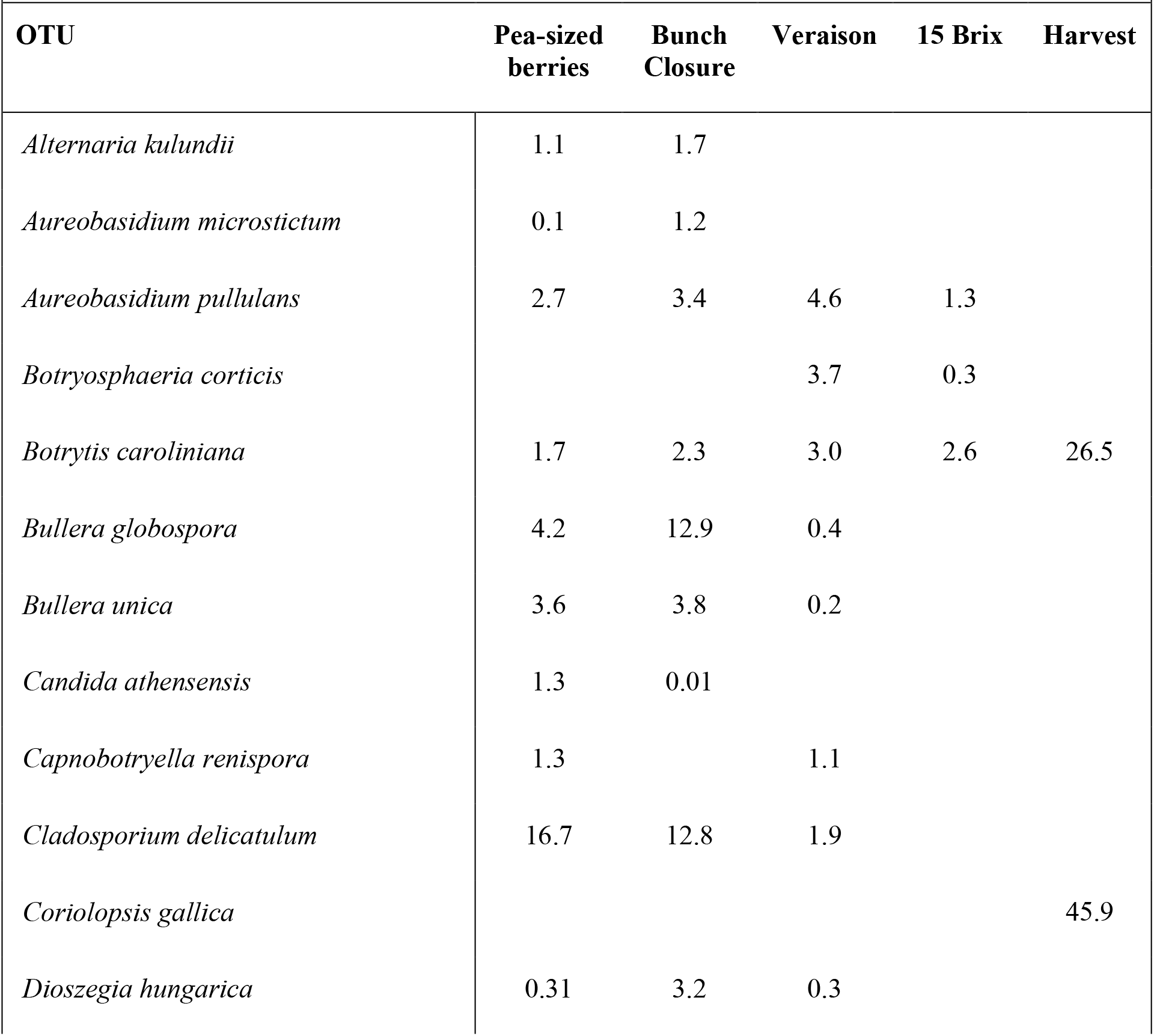
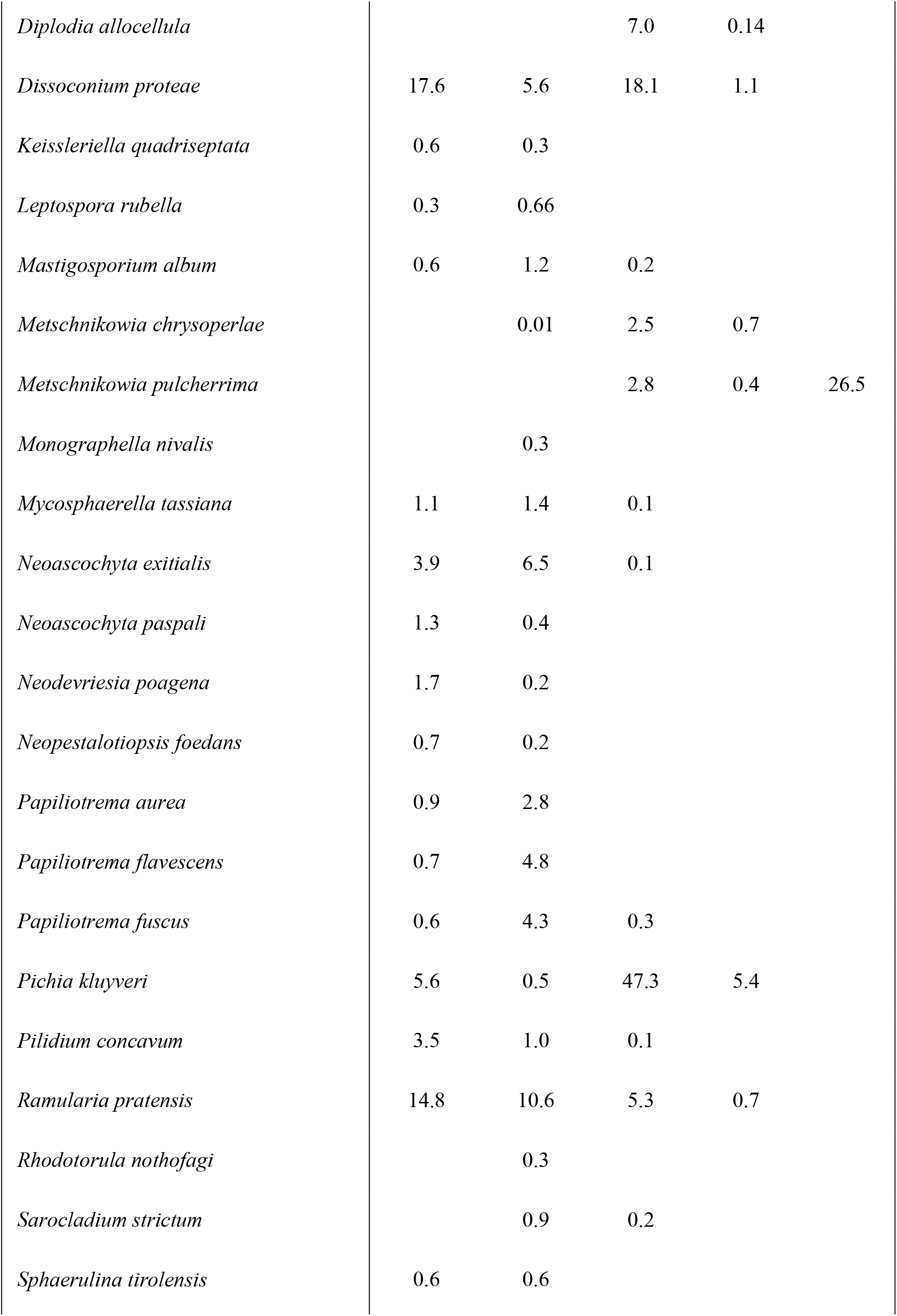
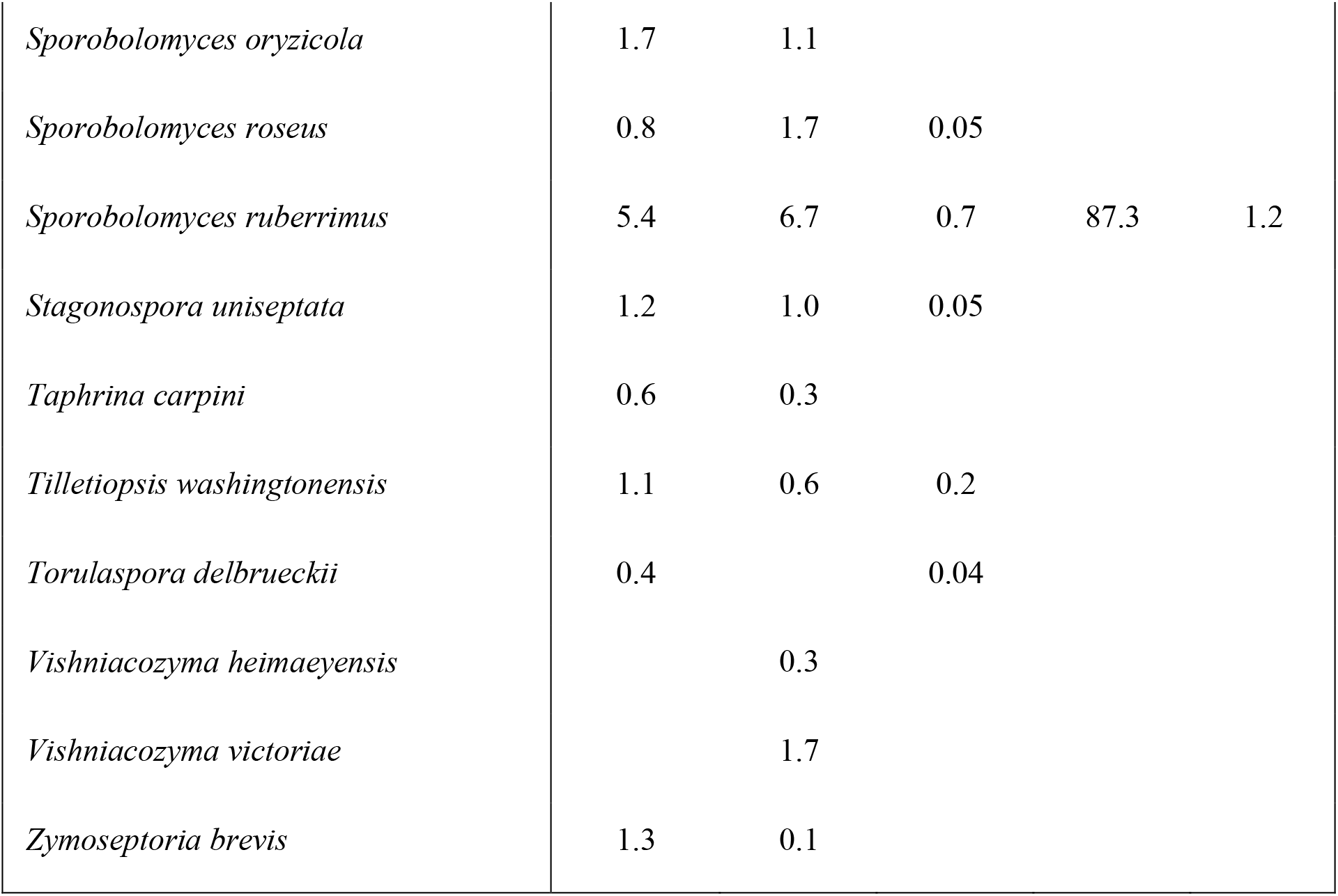
In 2015 Finger Lakes, New York, the relative mean frequency (%) of reads for each fungal OTU across three vineyards at five phenological stages. Sample numbers per stage are presented in Table 1.

**Table 5.**
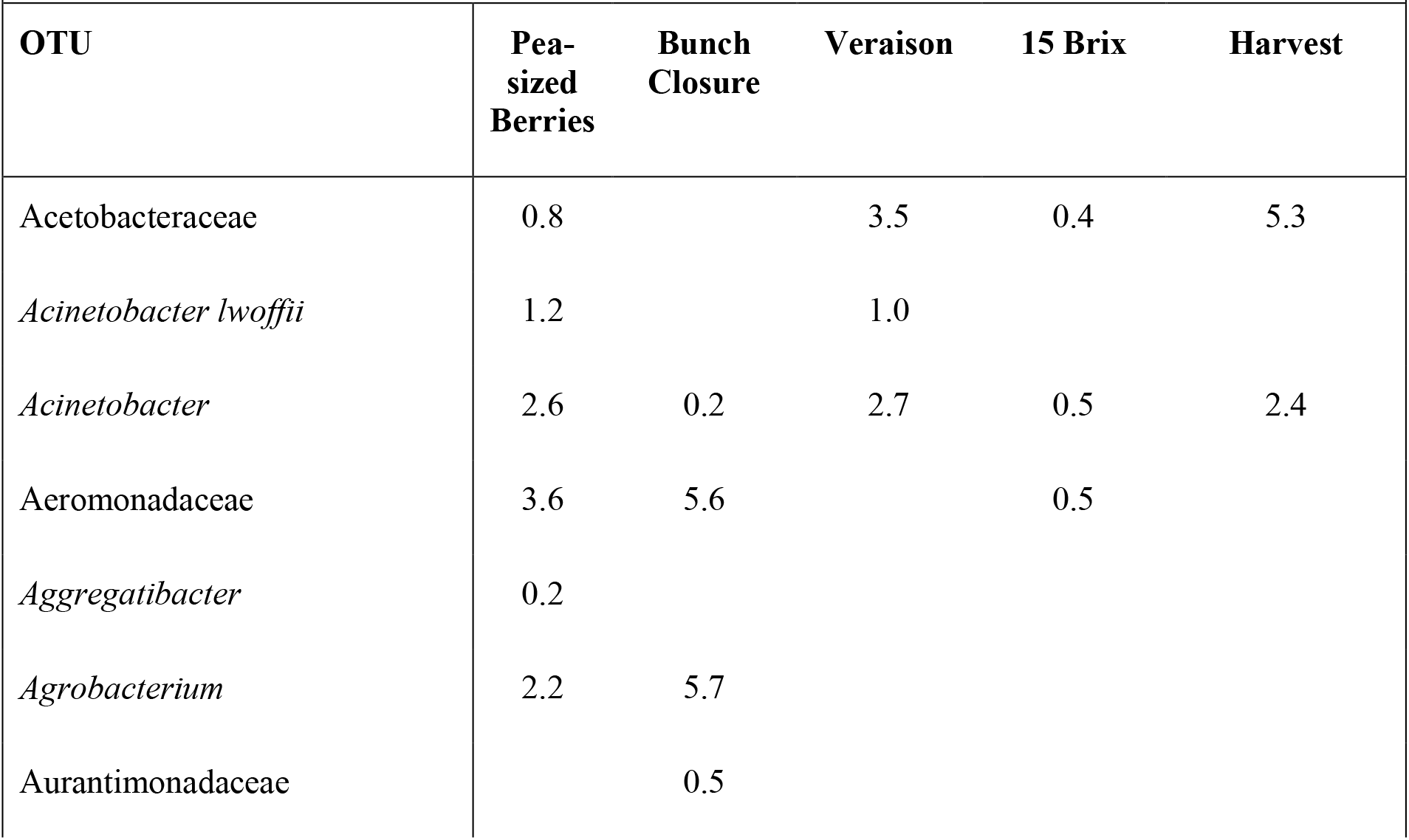
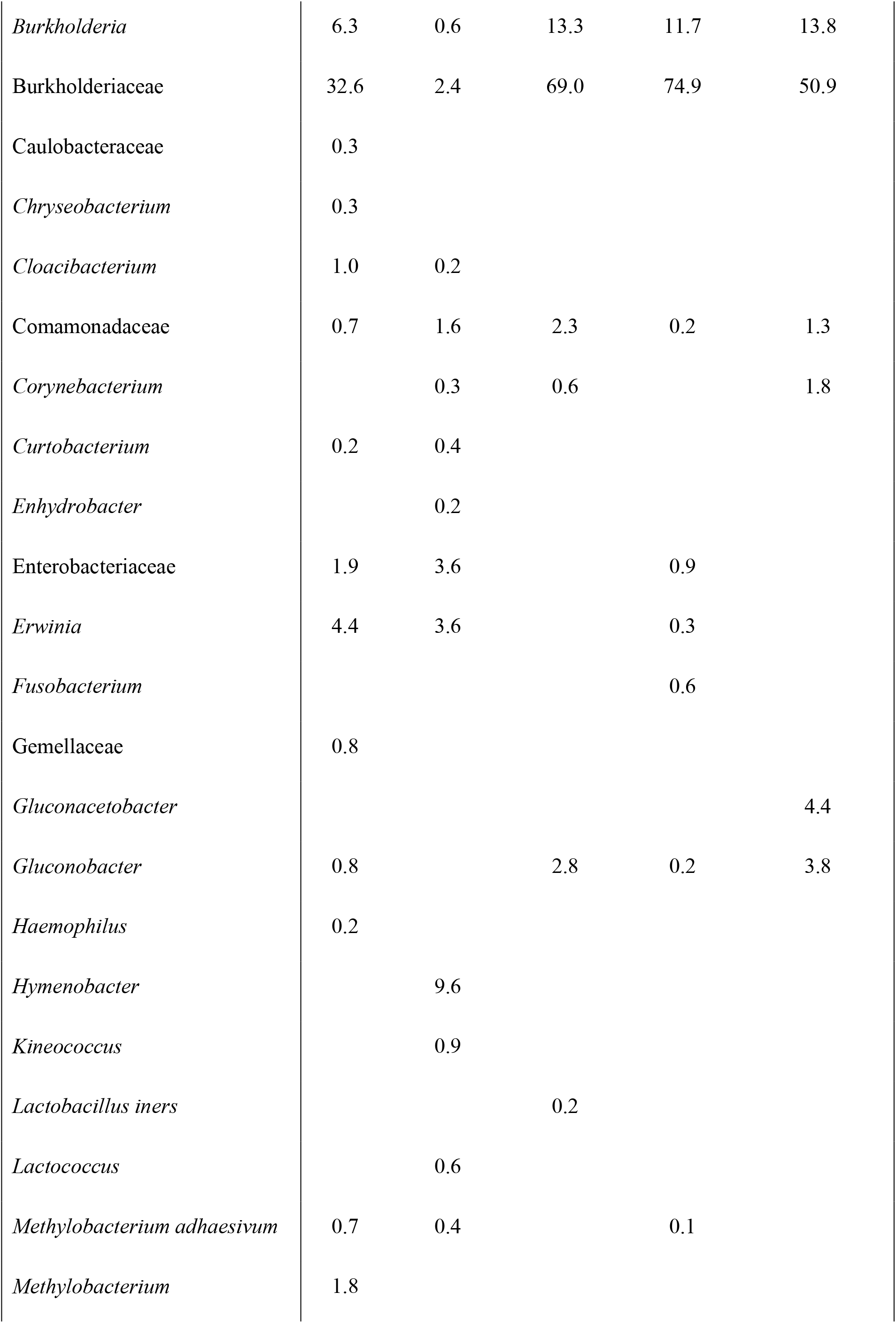
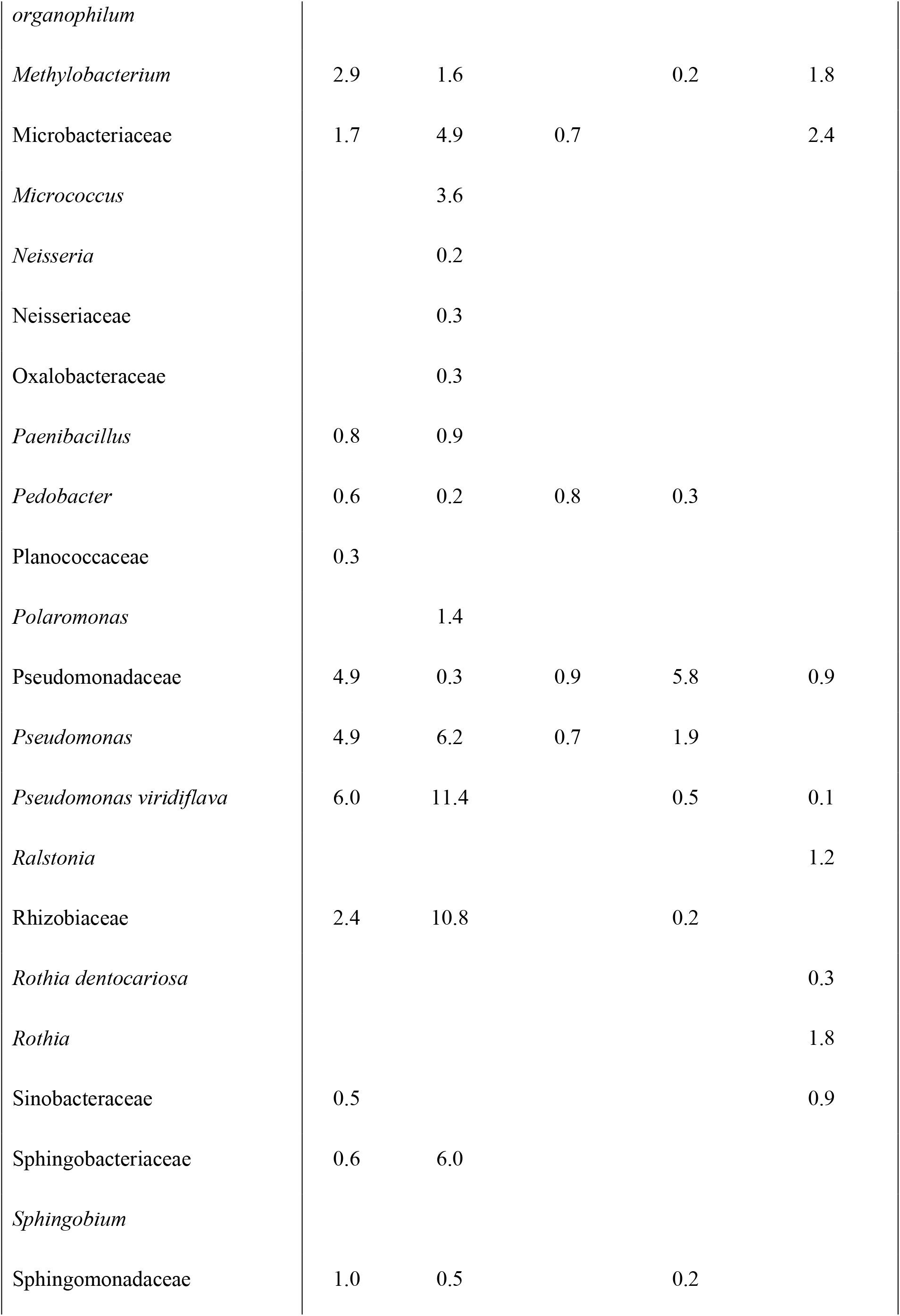
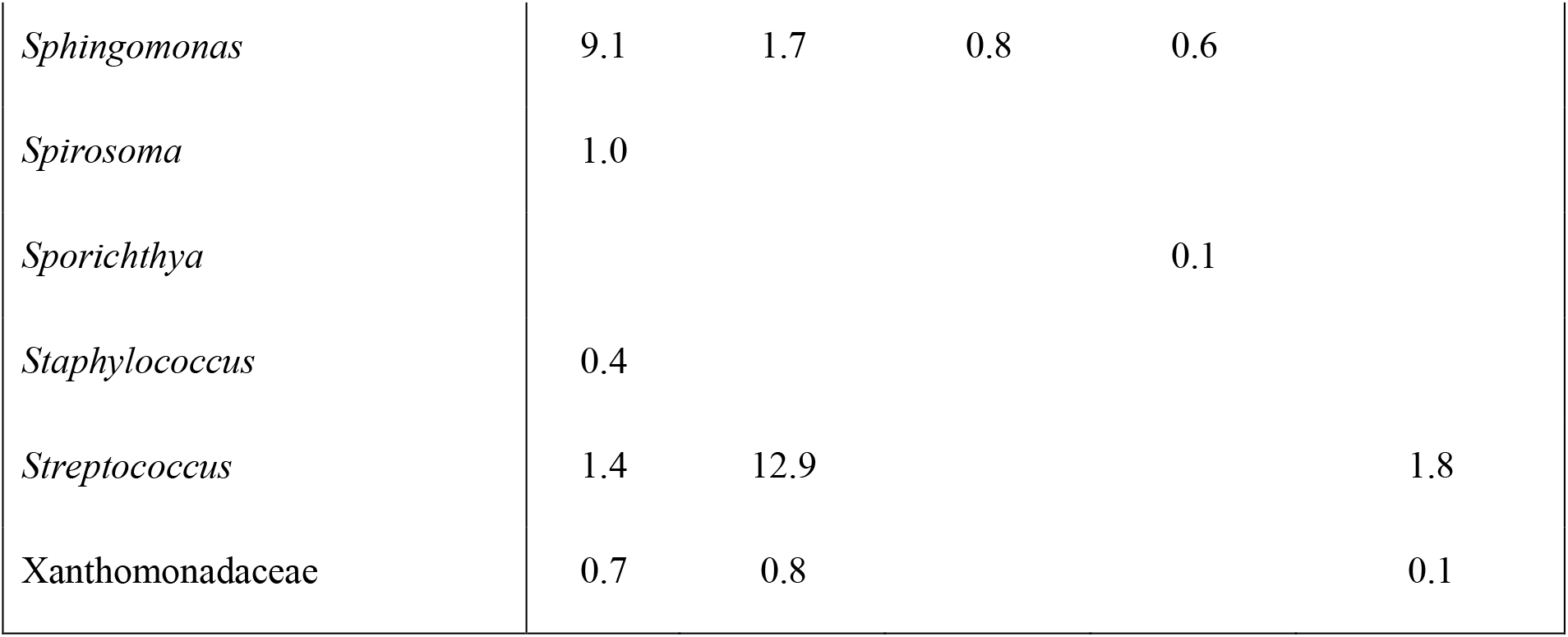
In 2015 Finger Lakes, New York, the relative mean frequency (%) of reads for each bacterial OTU across three vineyards at five phenological stages. Sample numbers per stage are presented in Table 1.

The number of fungal and bacterial OTUs in 2016 were also higher than in 2014 (36 and 57, respectively). Members of order Saccharomycetales were the most abundant fungal OTUs. At pea-sized berries, *Pichia* spp. represented 80% of the total reads, while *Candida xylopsoci* represented only 1%. At bunch closure, *P. kluyveri, P. membranifaciens* and *P. terricola* represented 42% of the total reads and *C. xylopsoci*, 47% of the total reads. *Pichia* spp. represented 73% of the total OTUs at Veraison, with *Candida* spp. and *Hanseniaspora* spp. representing 6% and 8% respectively. At 15 Brix, *Pichia* species represented 70% of the total reads, and 77% of the total reads at harvest. The majority of OTUs were identified less than 1% of the time (Table 6). For bacterial OTUs, members of order Rhodospirillales dominated every time point. *Gluconobacter* represented 22% of reads at pea-sized berries, 42% at bunch closure, 59% at Veraison, 23% at 15 Brix and 15% at harvest, and it also represented a significant proportion of reads from order Aceteobacteraceae. Members of the order Bacillaceae represented 37% of the OTUs are pea-sized berries, 10% at bunch closure, 20% at Veraison, 35% at 15 Brix and 44% at harvest (Table 7).

**Table 6.**
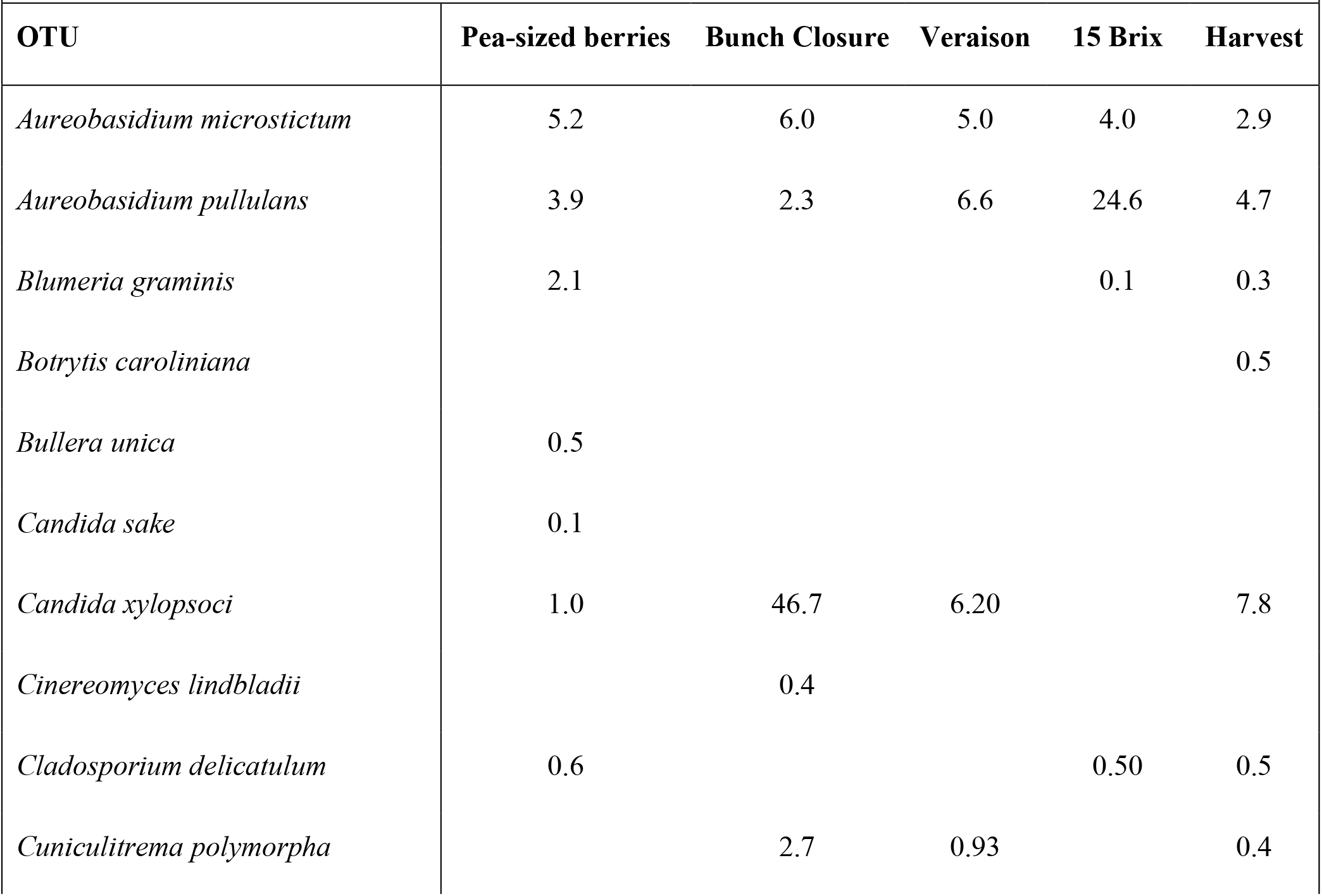
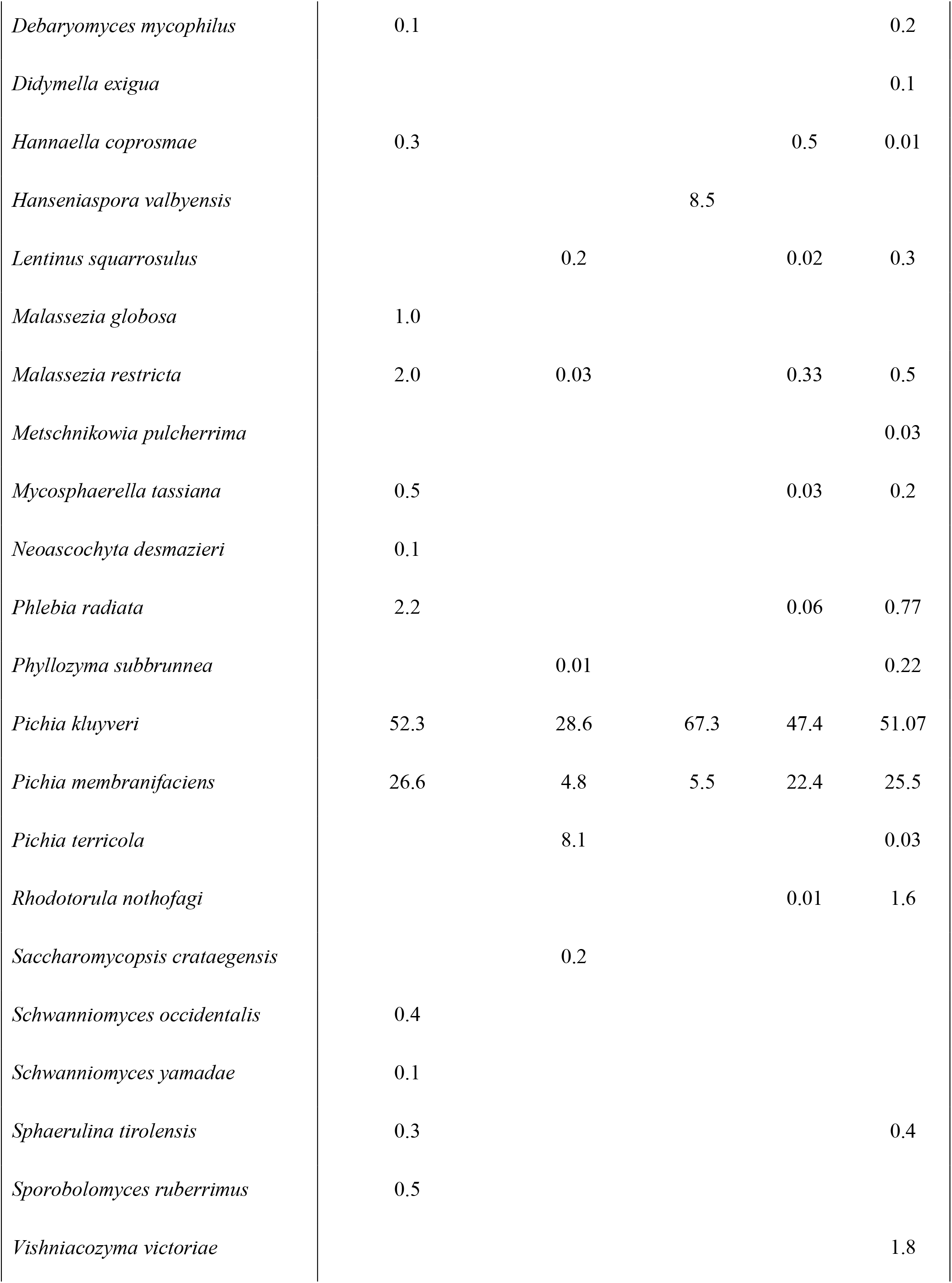
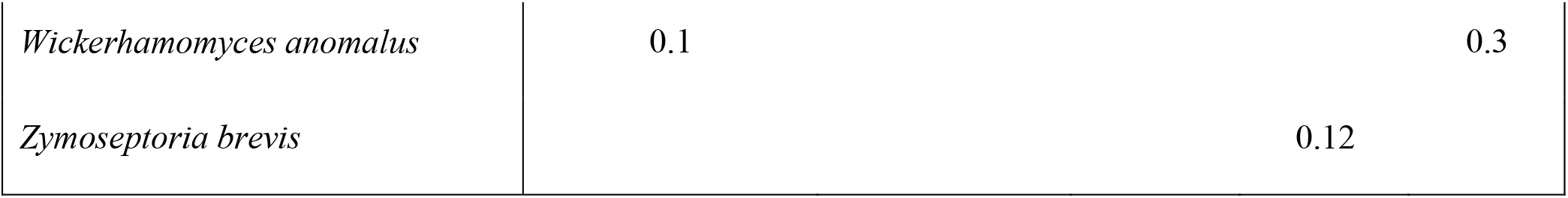
In 2016 Tasmania, Australia, the relative mean frequency (%) of reads for each fungal OTU across three vineyards at five phenological stages. Sample numbers per stage are presented in Table 1.

**Table 7.**
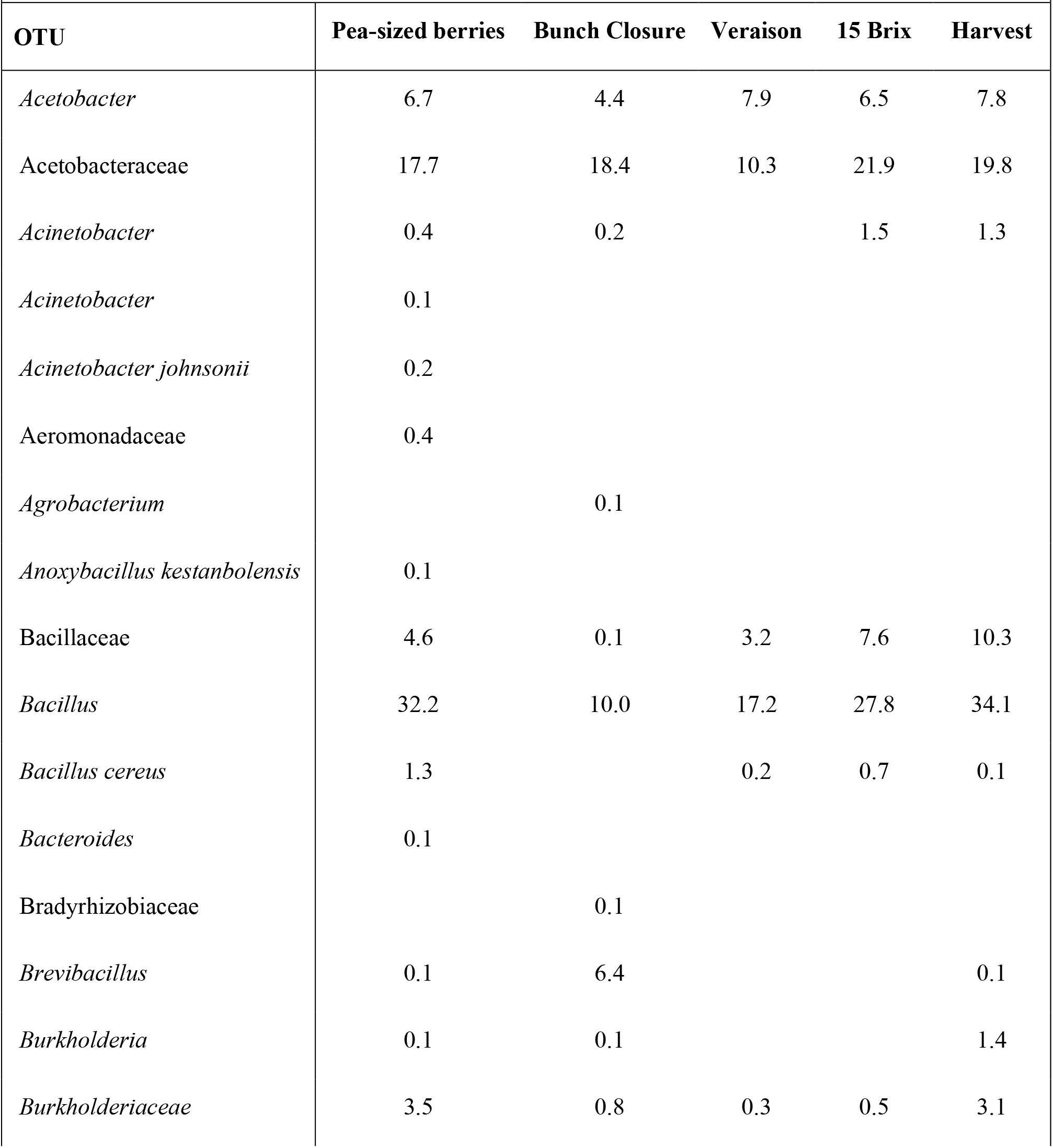
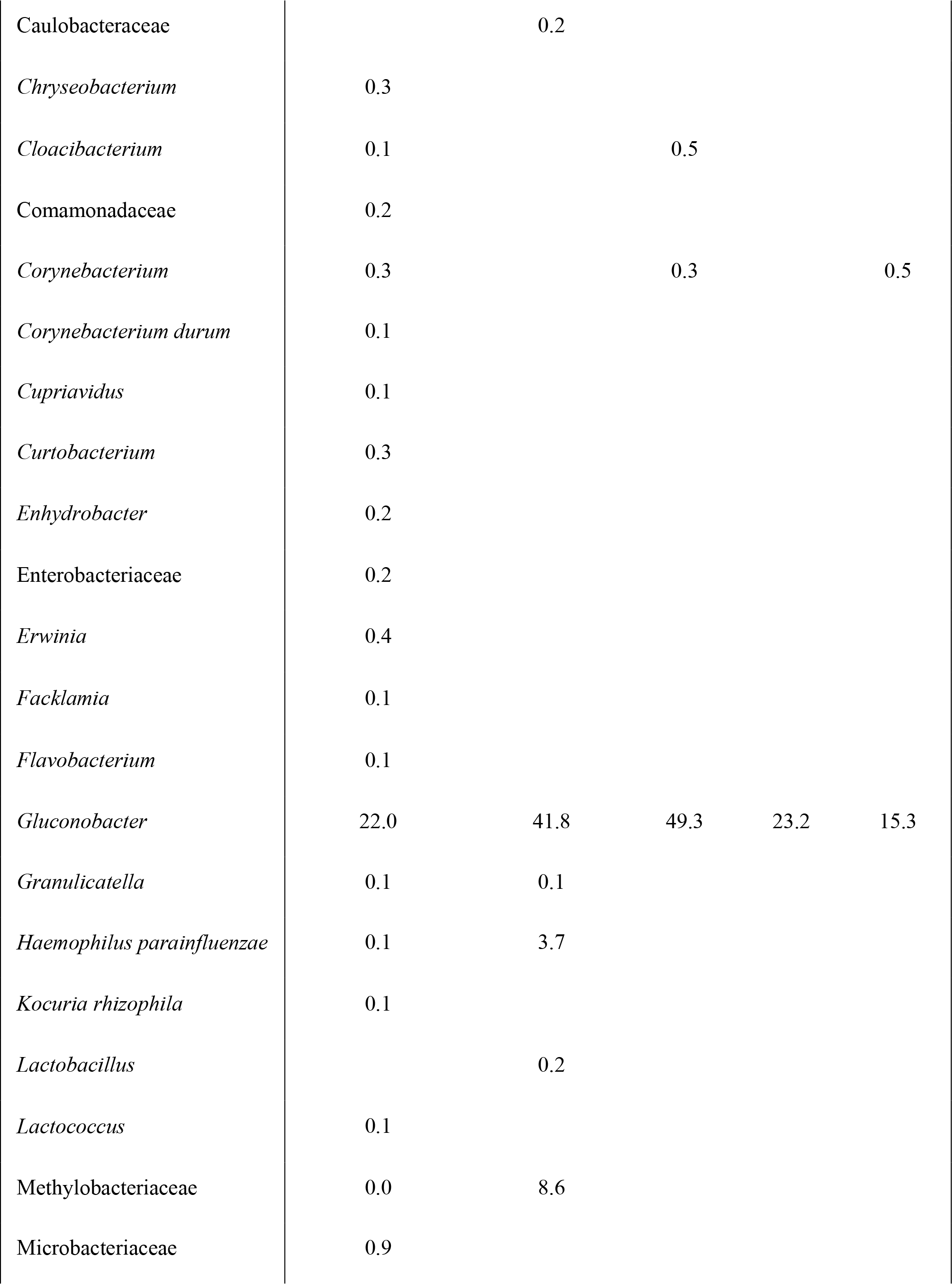
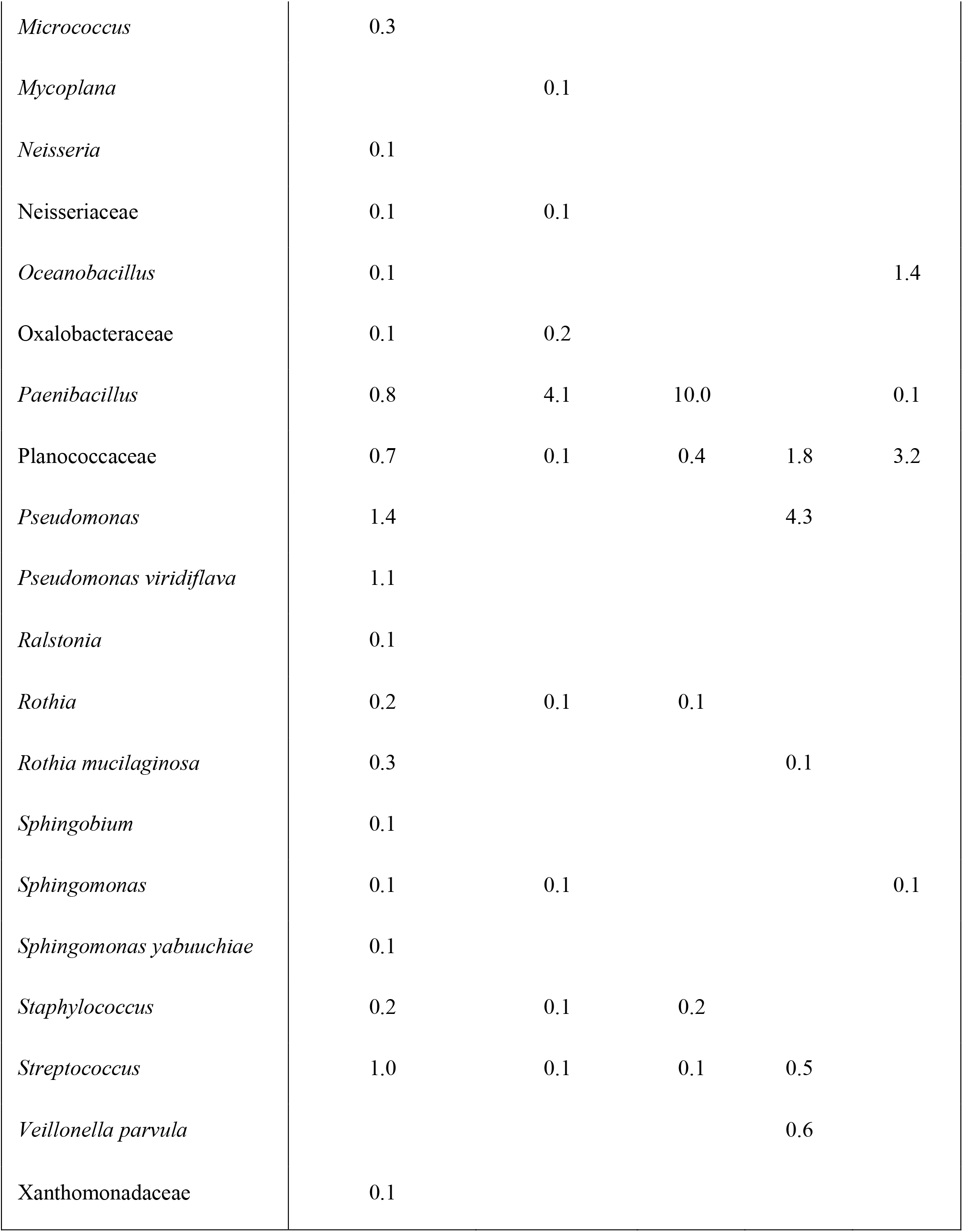
In 2016 Tasmania, Australia, the relative mean frequency (%) of reads for each bacterial OTU across three vineyards at five phenological stages. Sample numbers per stage are presented in Table 1.

|  |  |  |  |  |  |
| --- | --- | --- | --- | --- | --- |
| **OTU** | **Pea-sized berries** | **Bunch Closure** | **Veraison** | **15 Brix** | **Harvest** |
| *Acetobacter* | 6.7 | 4.4 | 7.9 | 6.5 | 7.8 |
| Acetobacteraceae | 17.7 | 18.4 | 10.3 | 21.9 | 19.8 |
| *Acinetobacter* | 0.4 | 0.2 |  | 1.5 | 1.3 |
| *Acinetobacter* | 0.1 |  |  |  |  |
| *Acinetobacter johnsonii* | 0.2 |  |  |  |  |
| Aeromonadaceae | 0.4 |  |  |  |  |
| *Agrobacterium* |  | 0.1 |  |  |  |
| *Anoxybacillus kestanbolensis* | 0.1 |  |  |  |  |
| Bacillaceae | 4.6 | 0.1 | 3.2 | 7.6 | 10.3 |
| *Bacillus* | 32.2 | 10.0 | 17.2 | 27.8 | 34.1 |
| *Bacillus cereus* | 1.3 |  | 0.2 | 0.7 | 0.1 |
| *Bacteroides* | 0.1 |  |  |  |  |
| Bradyrhizobiaceae |  | 0.1 |  |  |  |
| *Brevibacillus* | 0.1 | 6.4 |  |  | 0.1 |
| *Burkholderia* | 0.1 | 0.1 |  |  | 1.4 |
| *Burkholderiaceae* | 3.5 | 0.8 | 0.3 | 0.5 | 3.1 |
| Caulobacteraceae |  | 0.2 |  |  |  |
| *Chryseobacterium* | 0.3 |  |  |  |  |
| *Cloacibacterium* | 0.1 |  | 0.5 |  |  |
| Comamonadaceae | 0.2 |  |  |  |  |
| *Corynebacterium* | 0.3 |  | 0.3 |  | 0.5 |
| *Corynebacterium durum* | 0.1 |  |  |  |  |
| *Cupriavidus* | 0.1 |  |  |  |  |
| *Curtobacterium* | 0.3 |  |  |  |  |
| *Enhydrobacter* | 0.2 |  |  |  |  |
| Enterobacteriaceae | 0.2 |  |  |  |  |
| *Erwinia* | 0.4 |  |  |  |  |
| *Facklamia* | 0.1 |  |  |  |  |
| *Flavobacterium* | 0.1 |  |  |  |  |
| *Gluconobacter* | 22.0 | 41.8 | 49.3 | 23.2 | 15.3 |
| *Granulicatella* | 0.1 | 0.1 |  |  |  |
| *Haemophilus parainfluenzae* | 0.1 | 3.7 |  |  |  |
| *Kocuria rhizophila* | 0.1 |  |  |  |  |
| *Lactobacillus* |  | 0.2 |  |  |  |
| *Lactococcus* | 0.1 |  |  |  |  |
| Methylobacteriaceae | 0.0 | 8.6 |  |  |  |
| Microbacteriaceae | 0.9 |  |  |  |  |
| *Micrococcus* | 0.3 |  |  |  |  |
| *Mycoplana* |  | 0.1 |  |  |  |
| *Neisseria* | 0.1 |  |  |  |  |
| Neisseriaceae | 0.1 | 0.1 |  |  |  |
| *Oceanobacillus* | 0.1 |  |  |  | 1.4 |
| Oxalobacteraceae | 0.1 | 0.2 |  |  |  |
| *Paenibacillus* | 0.8 | 4.1 | 10.0 |  | 0.1 |
| Planococcaceae | 0.7 | 0.1 | 0.4 | 1.8 | 3.2 |
| *Pseudomonas* | 1.4 |  |  | 4.3 |  |
| *Pseudomonas viridiflava* | 1.1 |  |  |  |  |
| *Ralstonia* | 0.1 |  |  |  |  |
| *Rothia* | 0.2 | 0.1 | 0.1 |  |  |
| *Rothia mucilaginosa* | 0.3 |  |  | 0.1 |  |
| *Sphingobium* | 0.1 |  |  |  |  |
| *Sphingomonas* | 0.1 | 0.1 |  |  | 0.1 |
| *Sphingomonas yabuuchiae* | 0.1 |  |  |  |  |
| *Staphylococcus* | 0.2 | 0.1 | 0.2 |  |  |
| *Streptococcus* | 1.0 | 0.1 | 0.1 | 0.5 |  |
| *Veillonella parvula* |  |  |  | 0.6 |  |
| Xanthomonadaceae | 0.1 |  |  |  |  |

Heatmaps of the various years of the experiments by fungi and bacteria show that there is little pattern from one time point to the next (Figs. 1-5). There was not sufficient fungal data generated in 2014 to generate a heatmap. The heatmap of bacteria in 2014 show that 15 Brix most closely resembles the bunch closure timepoint, a trend also present in the 2015 bacteria (Figs. 1, 3). In the 2014 bacteria (Fig. 1) and 2016 fungi (Fig. 5), 15 Brix varied the most from the samples at harvest. Across all heatmaps, there is no pattern detected in terms of number of OTUs by time point or region. No time point was consistently low or high the number of bacterial or fungal OTUs in any year or region.

**Figure 1.**
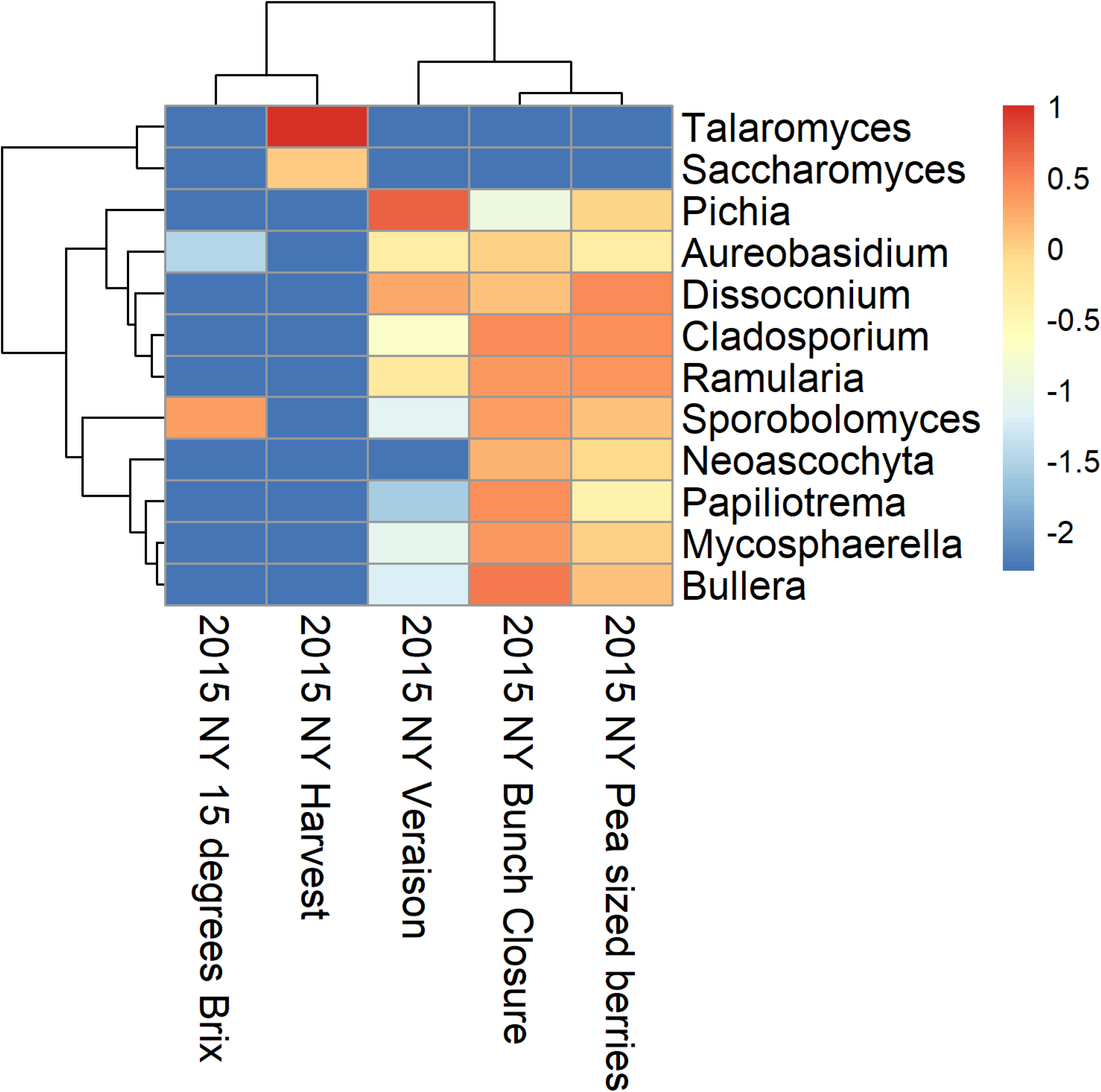
Heatmap representing 131 bacterial samples collected in 2014 in the Finger Lakes region of New York across five phenological timepoints: pea-sized, bunch closure, Veraison, 15 Brix and harvest. Hierarchical clustering was conducted using the complete method. The rows were clustered using the Euclidean method, and the columns were clustered using the Manhattan method.

**Figure 2.**
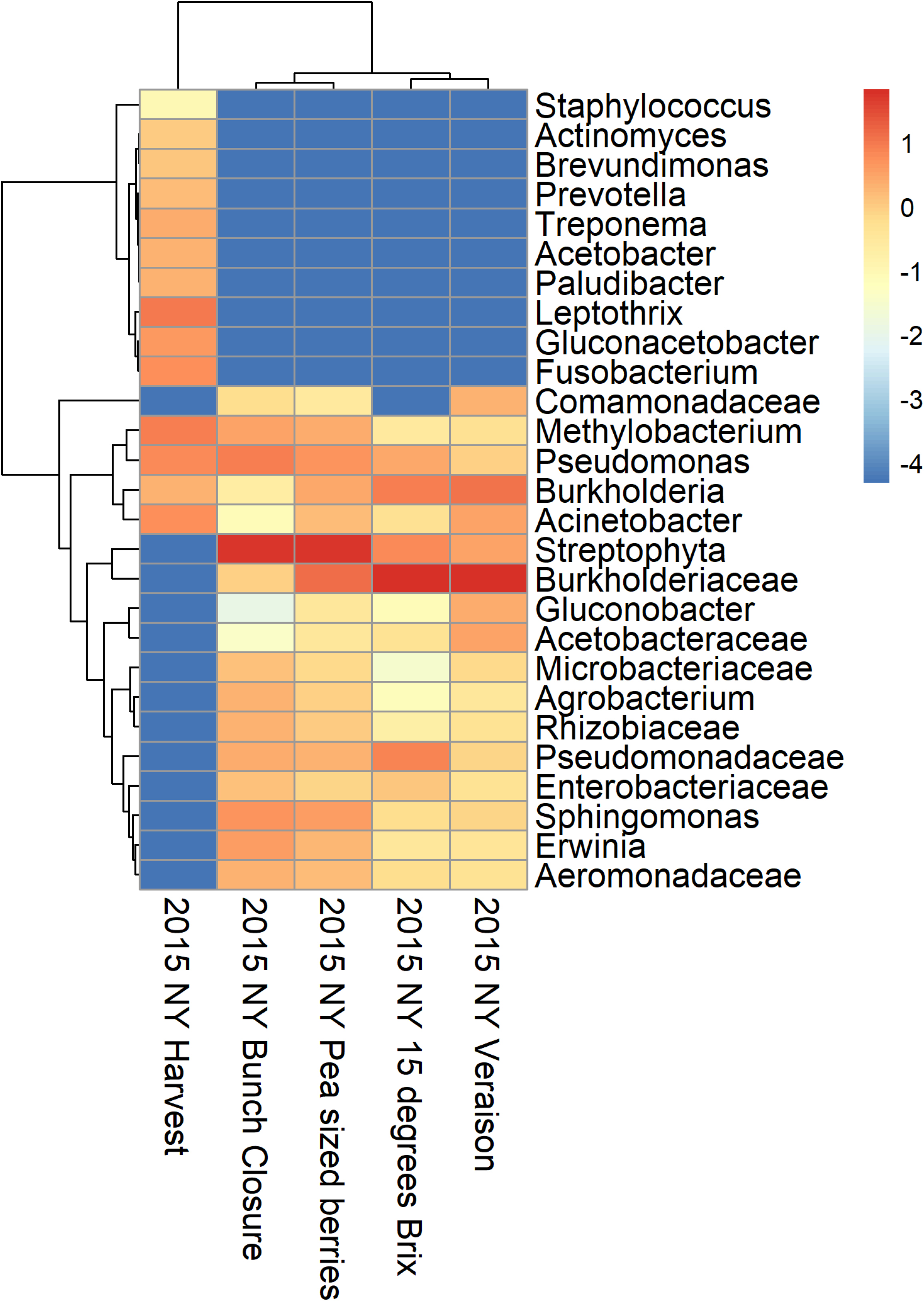
Heatmap representing 150 fungal samples collected in 2015 in the Finger Lakes region of New York across five phenological timepoints: pea-sized, bunch closure, Veraison, 15 Brix and harvest. Hierarchical clustering was conducted using the complete method. The rows were clustered using the Euclidean method, and the columns were clustered using the Manhattan method.

**Figure 3.**
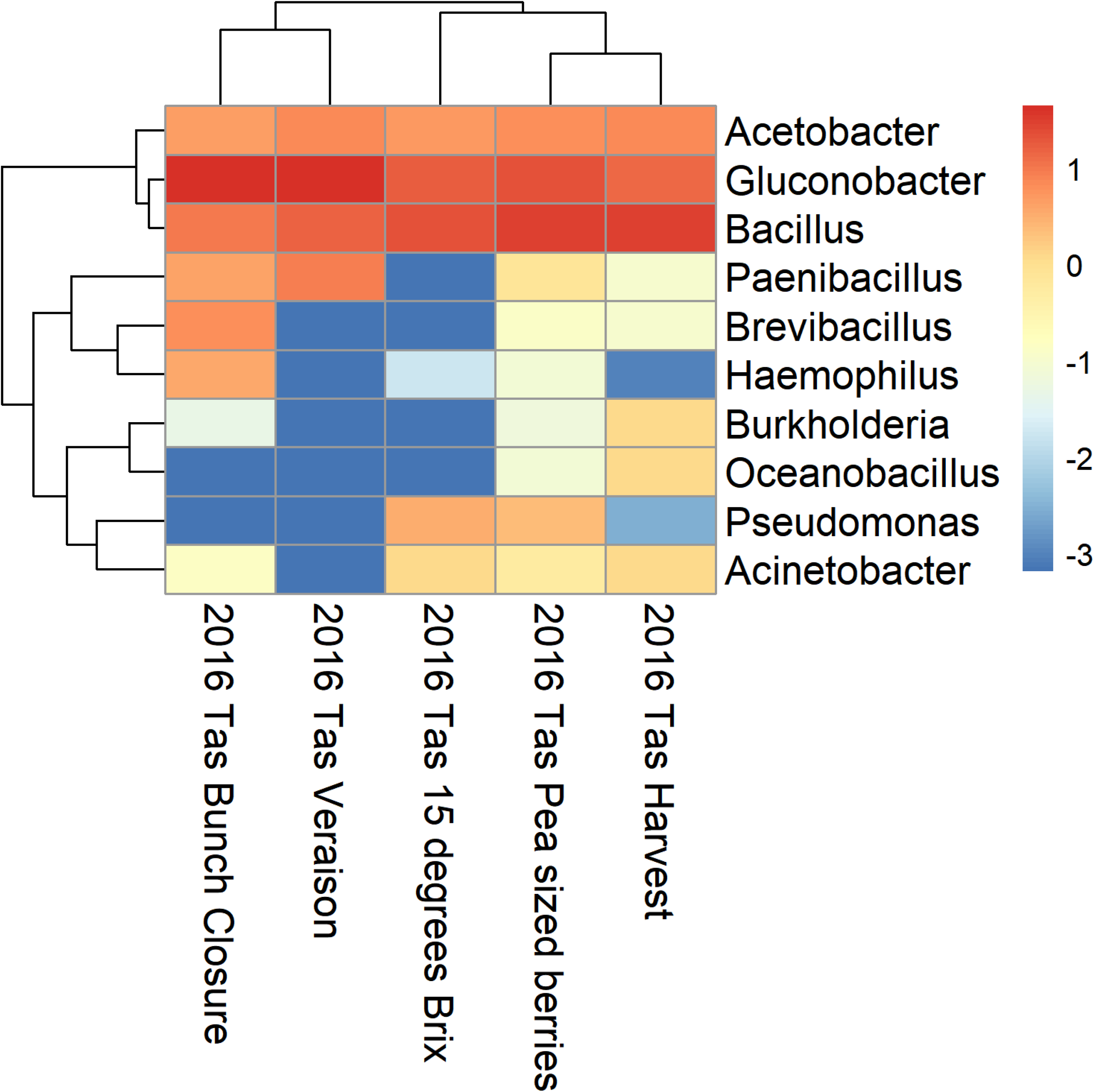
Heatmap representing 91 bacterial samples collected in 2015 in the Finger Lakes region of New York across five phenological timepoints: pea-sized, bunch closure, Veraison, 15 Brix and harvest. Hierarchical clustering was conducted using the complete method. The rows were clustered using the Euclidean method, and the columns were clustered using the Manhattan method.

**Figure 4.**
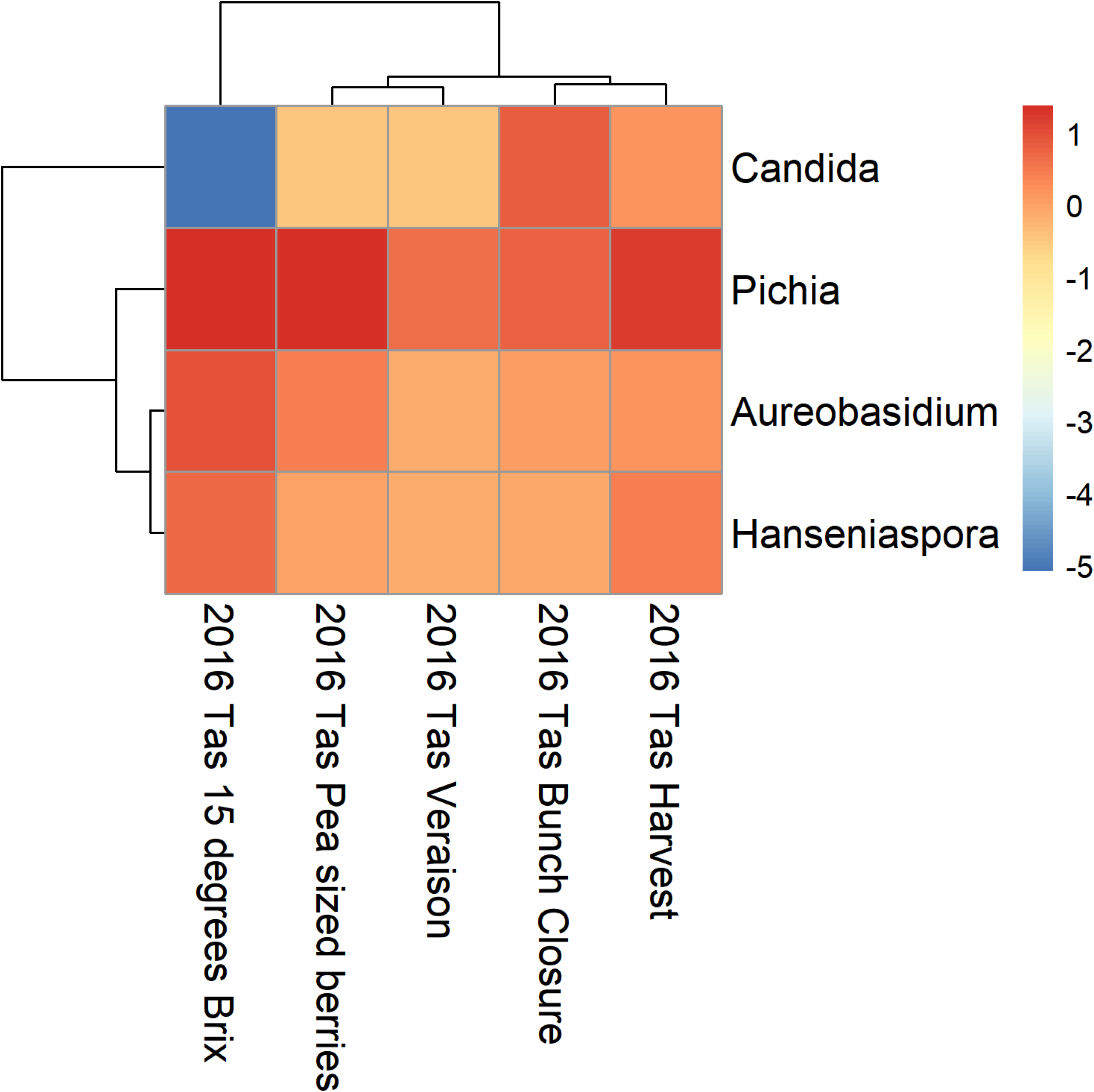
Heatmap representing 178 bacterial samples collected in 2016 in Tasmania, Australia across five phenological timepoints: pea-sized, bunch closure, Veraison, 15 Brix and harvest. Hierarchical clustering was conducted using the complete method. The rows were clustered using the Euclidean method, and the columns were clustered using the Manhattan method.

**Figure 5.**
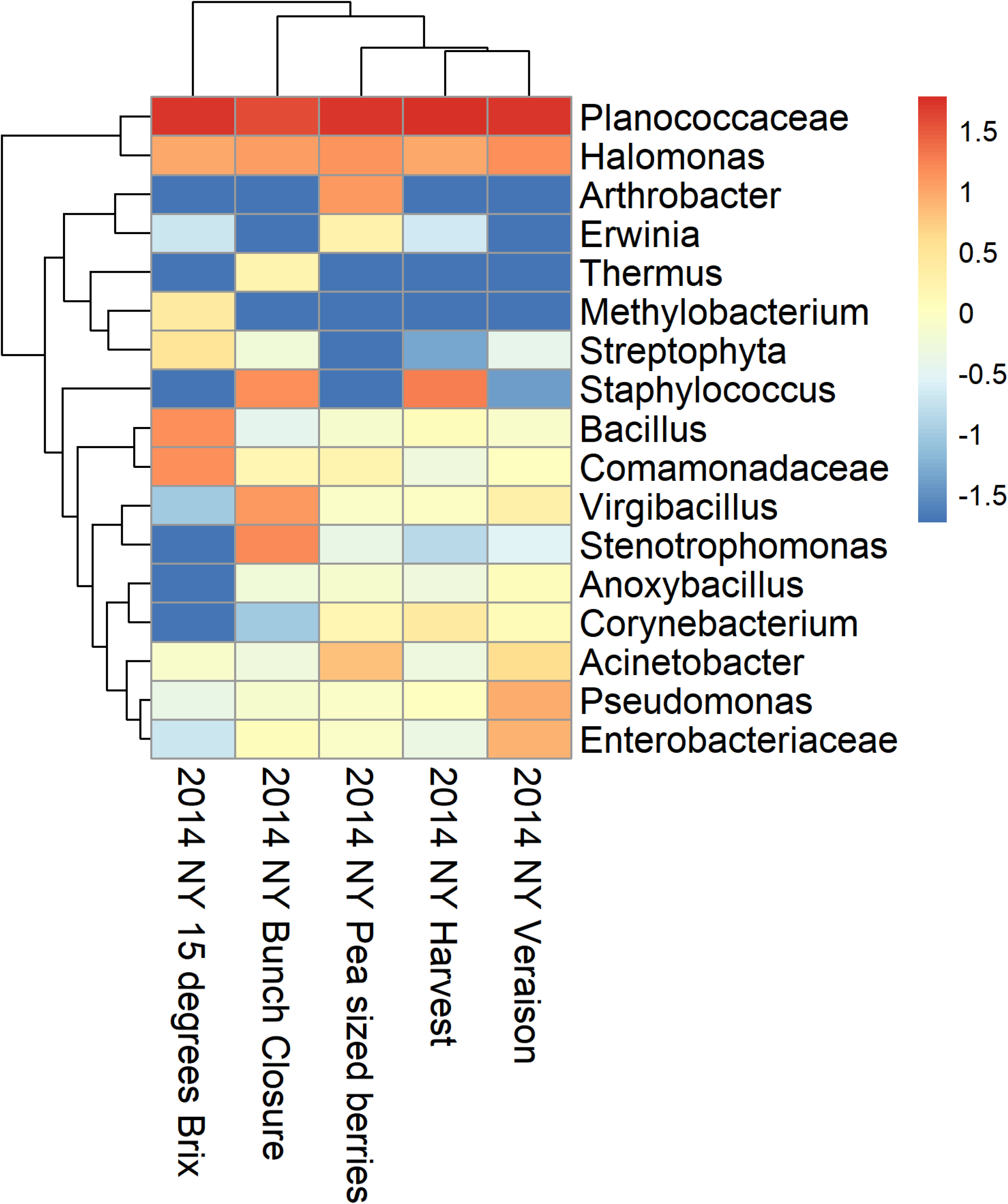
Heatmap representing 306 fungal samples collected in 2016 in Tasmania, Australia across five phenological timepoints: pea-sized, bunch closure, Veraison, 15 Brix and harvest. Hierarchical clustering was conducted using the complete method. The rows were clustered using the Euclidean method, and the columns were clustered using the Manhattan method.

## Discussion

The grape microbiota has become a popular subject of research in recent years, particularly with widespread adoption of high-throughput sequencing and metagenomics tools. While previous research focused primarily on microbial populations at harvest and in the grape must, our investigation explored the epiphytic population dynamics of grape microbiota season-long at key phenological stages over three years and two distinct grape growing regions.

On July 31, 2014, between bunch closure and Veraison, the Finger Lakes region suffered a major hail storm which severely impacted grape development, and it is after this event that we see a large and temporary spike in *Burkholderiaceae, Pichia kluyveri* and *Dissoconium proteae* and significant reduction in *Pseudomonas* spp., *Cladosporium delicatulum* and *Bullera globospora*. In 2015 in the Finger Lakes, there was a significant amount of sour rot near harvest (Hall et al., 2018b) and in Tasmania in 2016, the season was very dry but with significant sour rot infections near harvest (Hall and Wilcox, 2019a). The data from 2015 and 2016 has a larger representation of organisms at every time point than those data from 2014, along with a significantly higher percentage of yeast and acetic acid bacteria in the samples from 2015 and 2016, even as early as pea-sized berries. It is unknown whether the increased number of OTUs has an association with disease development or whether they are unrelated, because those microorganisms that play a role in the sour rot disease complex are also ubiquitous yeast and acetic acid bacteria on the grape surface (Hall et al., 2018a). The notable lack of those organisms in the 2014 data may be an indication of why sour rot infections were not prevalent that year, however.

There is a notable similarity between those data collected in 2015 and 2016, primarily in the increased number of microbial species, in comparison to the 2014 samples, and in the prevalence of yeast and acetic acid bacteria. Also significant are the differences between the 2014 and 2015 data. Since the data are from the same region, the differences were more significant than we expected. We recognize that in combining the results from many vineyards, we are not focusing on the microbial terroir of a single vineyard and how it changed from one year to the next, but examining the microbial terroir of a region allowed us to look at any patterns among multiple sites. However, because there are limited similarities between the microbial communities of 2014 and 2015 within the same sites, yet significant similarities between 2015 and 2016, despite being from different continents, it leads us to a much larger question how we describe microbial terroir. Researchers have examined microbial changes from different regions (Barata et al., 2012), and as our research indicates that the microbial terroir may in fact change dramatically from one year to the next.

The ebb and flow of organisms as the season progresses are an indication of how the environment may be impacting the growth of the grapes, or even how the microorganisms are responding to conventional sprays in the vineyard. Because we did not collect spray records for every vineyard from which we sampled, we cannot relate this data back to those specific applications. However, typical commercial production in these regions uses fungicide applications every 10 to 14 days with a rotation of chemistries, so certainly the sampled grapes were exposed to one or more sprays between each timepoint. The spike of *E. necator* reads in 2014 at 15 Brix is a possible example of how the population of that pathogen was controlled with a spray application. The increase of *Botrytis* spp. reads over the course of the season, however, could also be related to the sprays applied in those vineyards. And while this data gives us a broad look at the dynamics of the microbial system, further studies that relate microbial community data with spray applications would provide researchers with information about which microbes are being controlled with each spray, and which ones proliferate as a result of that population being controlled.

Grapes harbor a unique microbial community because those members have influence in the downstream processing of those grapes, especially as it relates to native fermentations. Researchers have focused on those microbes present at harvest, but these communities are changing and being influenced from the very start of the growing season. Through understanding how the dynamics of these microbial communities change over the course of the growing season, we can better understand how we arrive at the microbial communities that we encounter at harvest, and in the resulting grape must. Moreover, we can now examine more closely how spray applications throughout the season may influence the epiphytic microbiome, and how that relates to disease susceptibility throughout the growing season. While it is unclear how controlling for certain yeast or bacteria could influence the microbial community, it is also possible that counterbalancing the prevalence of certain organisms with those that are not pathogenic, could reduce the risk of disease symptoms development.

## Acknowledgments

The authors thank Finger Lakes and Tasmanian grape growers who allowed for sampling of their grapevines, as well as Katherine Evans at the University of Tasmania for coordination assistance. This work was supported the NY State Dept. of Agriculture and Markets, NY Wine and Grape Foundation, Specialty Crops Research Initiative and the Dyson Fund.

## Author Contributions

MH and LCD conceived and designed the experiments. WW secured funding for and consulted on sample collection and experimental design. MH performed the experiments and collected the data. IO performed the bioinformatics and MO supervised the bioinformatic analysis. MH wrote the manuscript.

## Importance

This is the first study, to the best of our knowledge, that examines the epiphytic microbiome of grapes across several time points during the growing season, and across several years and regions. Recent grape microbiome research has not distinguished between epiphytic and endophytic microorganisms, and has not focused on time points other than harvest. This research shows that the epiphytic microbiome of the grape is constantly changing throughout the growing season and is likely impacted by environmental factors as well as consistent chemical spray applications, which has implications for both vineyard management and winemaking practices.

